# Novel GVHD resistant humanized-PBMC mouse model for preclinical HIV research

**DOI:** 10.1101/2021.06.03.446835

**Authors:** Leo Holguin, Liliana Echavarria, John C. Burnett

## Abstract

Humanized mouse models are based on the engraftment of human cells in immunodeficient mouse strains, most notably the NSG strain. Most used models have a major limitation in common, the development of graft-versus-host disease (GVHD). GVHD not only introduces variabilities into the research data but also leads to animal welfare concerns. A new mouse strain, B6.129S-Rag2^tm1Fwa^ CD47^tm1Fpl^ Il2rg^tm1Wjl^/J which lacks Rag1, IL2rg, and CD47 (triple knockout or TKO), is resistant to GVHD development. We transplanted TKO mice with human peripheral blood mononuclear cells (PBMCs) to establish a new humanized PBMC (hu-PBMC) mouse model. A cohort of these mice was infected with HIV-1 and monitored for plasma HIV viremia and CD4^+^ T cell depletion. The onset and progression of GVHD were monitored by clinical signs. This study demonstrates that TKO mice transplanted with human PBMCs support engraftment of human immune cells in primary and secondary lymphoid tissues, rectum, and brain. Moreover, the TKO hu-PBMC model supports HIV-1 infection via intraperitoneal, rectal, or vaginal routes, as confirmed by robust plasma HIV viremia and CD4^+^ T cell depletion. Lastly, TKO mice showed a delayed onset of GVHD clinical signs (∼21 days) and exhibited significant decreases in plasma levels of TNFβ. Based on these results, the TKO hu-PBMC mouse model not only supports humanization and HIV-1 infection but is also resistant to GVHD development, making this model a valuable tool in HIV research.

**Importance:** Currently, there is no cure or vaccine for HIV infection, thus continued research is needed to end the HIV pandemic. While many animal models are used in HIV research, none is used more than the humanized mouse model. A major limitation with current humanized mouse models is the development of graft-versus-host disease (GVHD). Here, we show a novel humanized mouse model that is resistant to GVHD development and supports and models HIV infection comparable to well-established humanized mouse models.

## Introduction

Human Immunodeficiency Virus (HIV) is a global pandemic that currently affects 37 million people with about 2.3 million new cases every year (1). Currently, there is no effective cure or prophylactic vaccine for HIV infection therefore, continued research is needed to end the HIV pandemic. Animal models play an important role in finding new cures and vaccines. The use of humanized mice models has gained popularity due to their “humanized” immune system (2-4) and has been proven useful for the study of HIV replication, pathogenesis, transmission, and prevention. The most common humanized mice models for HIV research are engrafted with human peripheral mononuclear blood cells (hu-PBMC), CD34^+^ hematopoietic stem cells (hu-CD34^+^), or a combination of CD34^+^ cells and the implantation of fetal liver and thymic tissue (hu-BLT) (5). All of these models involve the use of immunodeficient mouse strains: NSG (NOD.Cg-Prkdc^scid^ Il2rg^tm1Wjl^ /Sz), NRG (NOD.129S7(B6)-Rag1^tm1Mom^ IL2rg^tm1Wjll^ /Sz), NOG (NOD.Cg-Prkdc^scid^ Il2rg^tm1Sug^), and BRG (C.129(Cg)Rag2^tm1Fwa^ IL2rg^tm1Sug^ /Jic); which can support efficient, stable and systemic engraftment of human cells and tissues (6).

The hu-PBMC model is a fast and logistically simple method for mouse humanization (5). Kim et al. described a simple method of transplanting human PBMC in non-irradiated NSG mice (7). In the NSG hu-PBMC model, CD45^+^, CD3^+^, CD4^+^, and CD8^+^ T cells were detected in peripheral blood, lymph nodes, spleen, and liver (7). The model supported HIV-1 infection with depletion of CD4^+^ T cell counts and high plasma viremia and responded to antiretroviral therapy (ART) with no detectable viremia or CD4^+^ T cell depletion (7). The hu-PBMC NSG model serves as a relevant HIV-1 infection and pathogenesis model that can be used in preclinical *in vivo* studies, however, it does have a major limitation.

A major limitation with the hu-PBMC model is the development of xenogeneic graft-versus-host disease (GVHD). In the hu-PBMC model, the onset of GVHD varies between 4-8 weeks after the transplantation of human PBMC (8). GVHD occurs when the engrafted human cells start to attack the mouse tissues leading to the early euthanasia of mice and loss of valuable data (9). Clinically, GVHD presents as weight loss, hunching posture, hair loss, and decreased activity/ lethargy (10). Besides limiting the experimental time of the animals, these clinical signs represent animal welfare concerns often in conflict with Institutional Animal Care and Use Committees (IACUC) standards. Pathologically, GVHD is characterized by lymphocytic infiltration and sclerosis of the skin, GI tract, and skin culminating in the early death of mice. Besides the loss of animals, GVHD can complicate the experimental results by introducing GVHD-associated fibroinflammatory lesions (11). A potential solution is the development of a GVHD-resistant hu-BLT model. Lavender et al. developed a hu-BLT model that develops B and T cell immunity to HIV infection and is resistant to GVHD, using the mouse strain B6.129S-Rag2^tm1Fwa^ CD47^tm1Fpl^ Il2rg^tm1Wjl^/J (TKO) (12). They observed that TKO hu-BLT mice did not develop any GVHD-related clinical signs over the 29 weeks of the experiment and in a subsequent study observed no GVHD development up to 45 weeks (13).TKO mice lack CD47 in addition to Rag 1 and IL2rg allowing for reconstitution of human immune cells with little GVHD development (12, 13).

In this study, we hypothesized that TKO mice can be used to develop a novel hu-PBMC model that has a delayed onset and reduced progression of GVHD and support HIV-1 infection. Here, we show that the TKO hu-PBMC model supports engraftment of human immune cells, supports HIV-1 infection, and showed a delayed presentation of GVHD clinical signs (∼21 days) compared to NSG mice. Based on these results, the TKO hu-PBMC mouse model not only supports humanization and HIV-1 infection but is also resistant to GVHD development, making this model a valuable tool in HIV research.

## Results

### TKO mice support the engraftment of human peripheral blood mononuclear cells

To test whether TKO mice were able to support engraftment of human PBMC, we engrafted 4-6 week old TKO mice with 10 million hu-PBMCs by IP injection as previously described (7). To compare the TKO mice with a standard PBMC model, we engrafted in parallel 4-6-week-old NSG mice with the same donor hu-PBMCs and analyzed peripheral blood every two weeks after transplantation. TKO mice showed detectable and stable circulating human CD45^+^ leukocytes starting at two weeks after transplantation comparable to the NSG mice (Fig. 1a). The majority of the CD45^+^ cells in the peripheral blood were CD3^+^ lymphocytes, which is also seen in the NSG hu-PBMC model (Fig. 1b). Further analysis of the CD3^+^ cells showed that TKO mice support robust circulating CD4^+^ T cells (Fig. 1c) and CD8^+^ T cells (Fig. 1d), similar to the NSG mice. In TKO mice, CD4^+^ T cells peaked at around week 6 and remained relatively stable over the 11-week study. In comparison, NSG mice reached peak CD4^+^ T cells around week 8 (Day 56) having a delay on peak circulating cells (Fig. 1c). This shows that the TKO hu-PBMC model can reliably engraft hu-PBMCs and repopulate CD3^+^ lymphocytes, including HIV target cells: CD4^+^ T cells.

**Figure 1.**
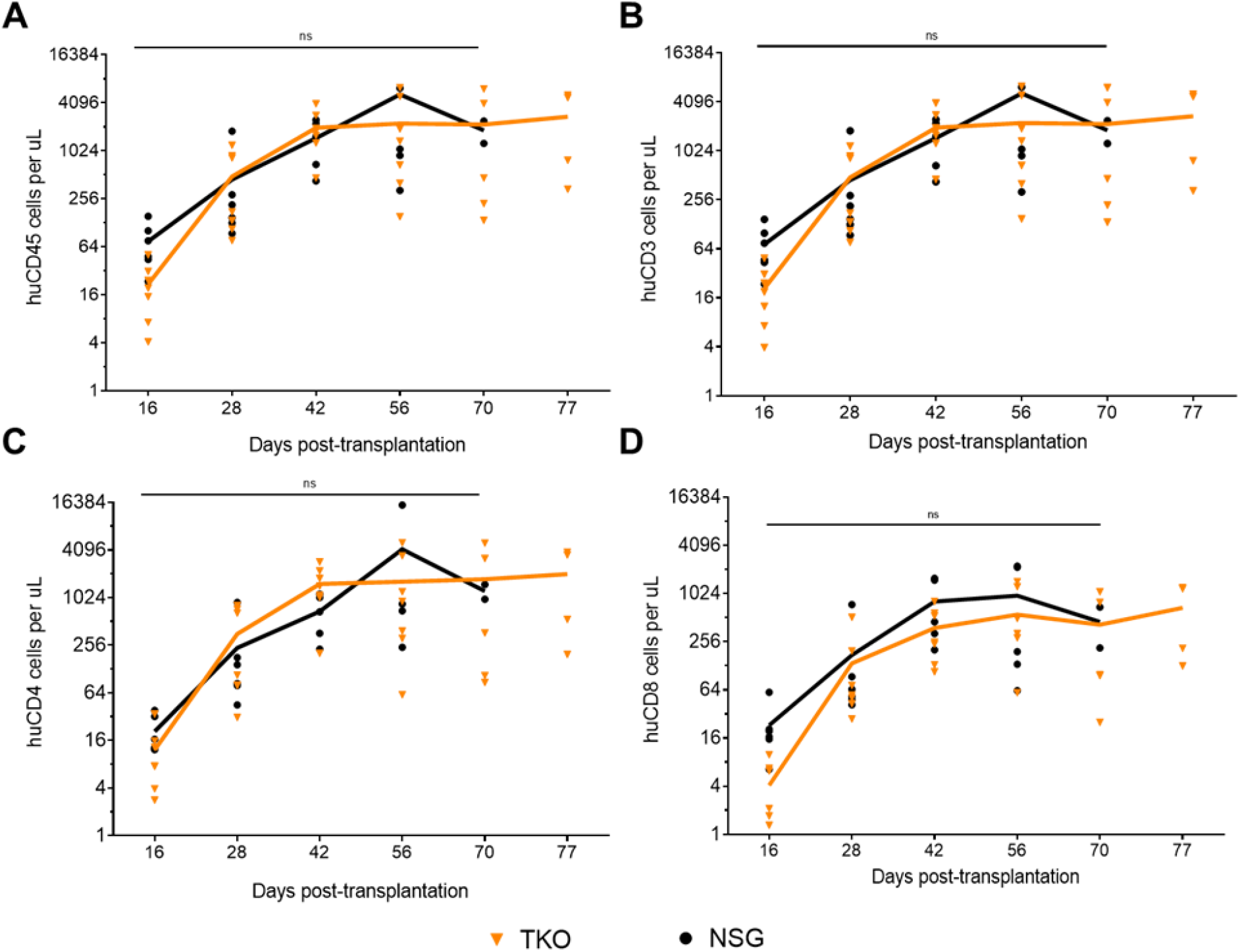
TKO mice support engraftment of human PBMCs comparable to NSG mice. 4-6-week-old TKO (n=7) and NSG (n=6) mice were transplanted with 1.0 × 10^7^ human PBMCs. Engraftment of human cells was checked by FACS analysis every 2 weeks for 11 weeks as shown in the graphs. Absolute cell numbers were calculated by using BD Liquid Counting Beads. Similar to the NSG hu-PBMC mice, TKO mice support engraftment of CD45^+^ leukocytes (**A**), mainly CD3^+^ T cells (**B**) including robust numbers of CD4^+^ (**C**) and CD8^+^ T cells (**D**). (3 TKO mice were lost before experiment endpoint at day 70 and 77, while all 6 NSG mice were lost before experiment endpoint at day 42, 70, and 77). Statistical significance was determined by unpaired t-test using the Holm-Sidak method on Prism, with alpha = 0.05.

Next, we analyzed the development of lymphoid tissues in the mice. Lymphoid tissues are sites where abundant lymphocytes live and sites of antigen presentation and lymphocyte activation in humans, which is important for HIV-1 replication and maintenance of persistent HIV-1 infection (14). After the eleven-week study, mice were humanely euthanized, and tissues were collected for immunohistochemistry analysis. TKO mice showed robust staining of CD3^+^ lymphocytes, CD4^+^ and CD8^+^ T cells in many tissues including bone marrow, lung, liver, rectum, and spleen, while CD14^+^/163^+^ macrophages and CD20^+^ B cells were mainly seen in lymph node and spleen (Fig. 2, Table, and Supplemental Fig. 1). Importantly, the spleen and bone marrow showed robust staining for CD4^+^ T cells indicating that the TKO hu-PBMC model supports the development of target HIV lymphoid tissue reservoirs. Collectively, the circulating CD4^+^ T cells (Fig. 1) and development of CD4^+^ T cell-rich lymphoid tissues (Fig. 2) suggest that the TKO hu-PBMC model may facilitate *in vivo* studies of HIV-1 infection.

**Figure 2.**
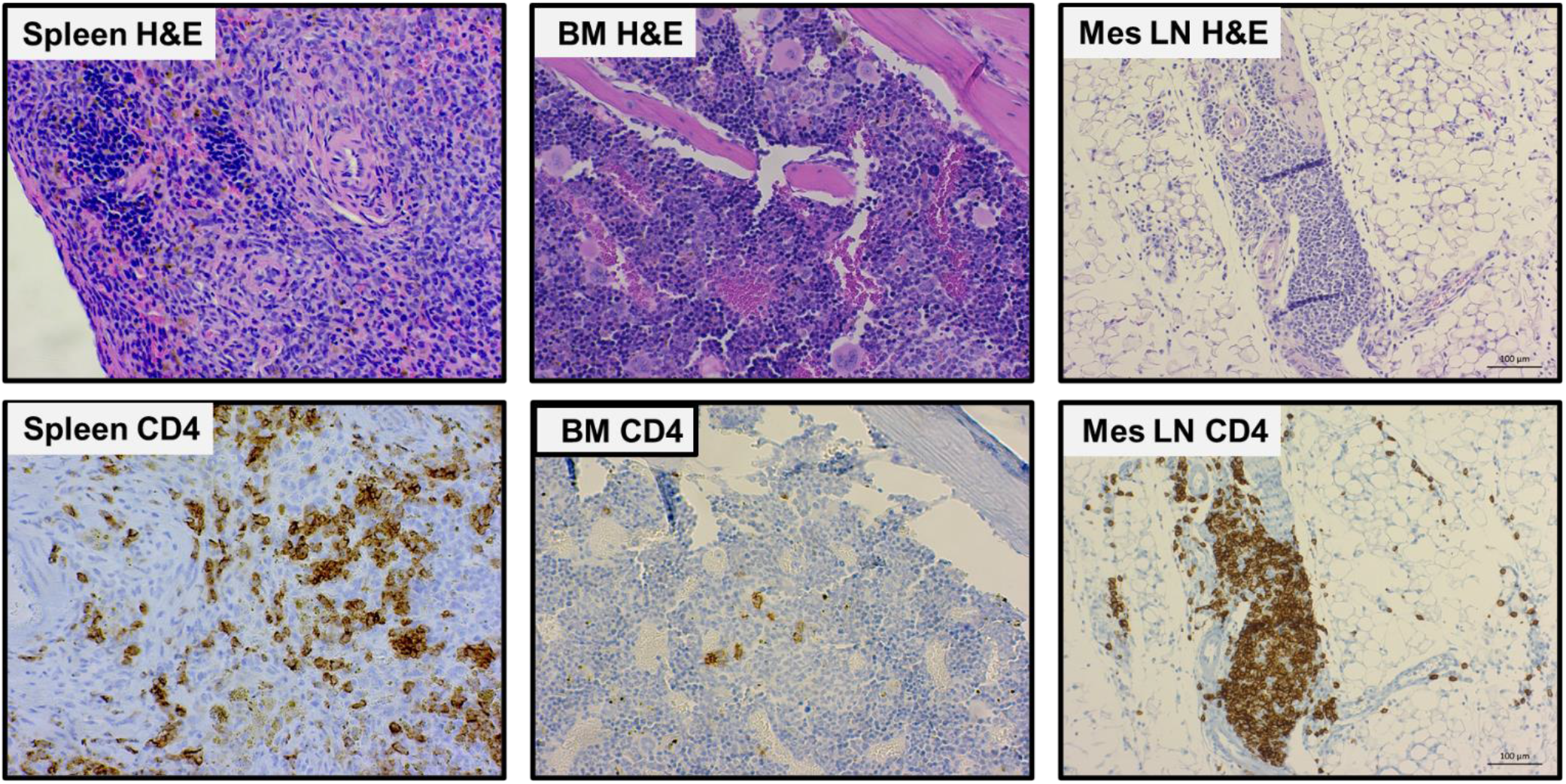
TKO hu-PBMC model supports engraftment of human T cells in lymphoid tissues. TKO mice with robust and consistent peripheral CD45^+^ engraftment were humanely euthanized, and tissues were collected for histology and IHC. Tissues were processed by the COH Solid Tumor Pathology Core for histology and immunohistochemistry: H&E, huCD3^+^, huCD4^+^, huCD8^+^. There is robust positive staining of CD3^+^, CD4^+^ (as seen above), and CD8^+^, supporting that the TKO hu-PBMC model supports engraftment of T cells in lymphoid tissues. Representative images are shown.

### TKO hu-PBMC model supports and models HIV-1 infection

To test whether the TKO hu-PBMC model supports HIV-1 infection, we challenged engrafted TKO mice with HIV-1BaL virus. About two weeks after transplantation, mice were bled and FACS analysis was done to confirm engraftment of mice. Once confirmed, mice were challenged with 200 ng p24 HIV-1BaL by IP injection under general anesthesia. NSG hu-PBMC mice were also challenged in parallel to compare the TKO model with the NSG model. After HIV challenge, plasma viremia and cell composition were monitored every two weeks by plasma qPCR and FACS analysis. TKO hu-PBMC mice began to show robust plasma viremia two weeks after HIV challenge and plasma viremia levels were maintained throughout the study with no significant differences to NSG hu-PBMC mice (Fig. 3a). We also monitored CD4^+^ T cell counts to see whether the hu-mouse model reflected depletion of CD4^+^ T cells seen in humans. Starting two weeks post-HIV challenge, CD4^+^ T cell counts begin to decline in both HIV infected TKO and NSG mice compared to control TKO mice (Fig. 3b). CD4^+^ T cell counts continue to decline over the remainder of the study. The data show that as plasma viral load increases in the mice, CD4^+^ T cells decrease over time, which accurately models HIV-1 infection in humans. To test the utility of the model, we challenged mice with HIV-1BaL via rectal and vaginal routes. Similar to intraperitoneal challenge, mice had robust plasma viremia and CD4^+^ T cell depletion when challenged rectally or vaginally with HIV-1 (Supplemental Fig. 2), although it was delayed compared to intraperitoneal challenge. These results show that TKO hu-PBMC mice challenged intraperitoneally, rectally, or vaginally with HIV-1 are readily infected with HIV-1 leading to robust plasma viremia and depletion of circulating CD4^+^ T cells.

**Figure 3.**
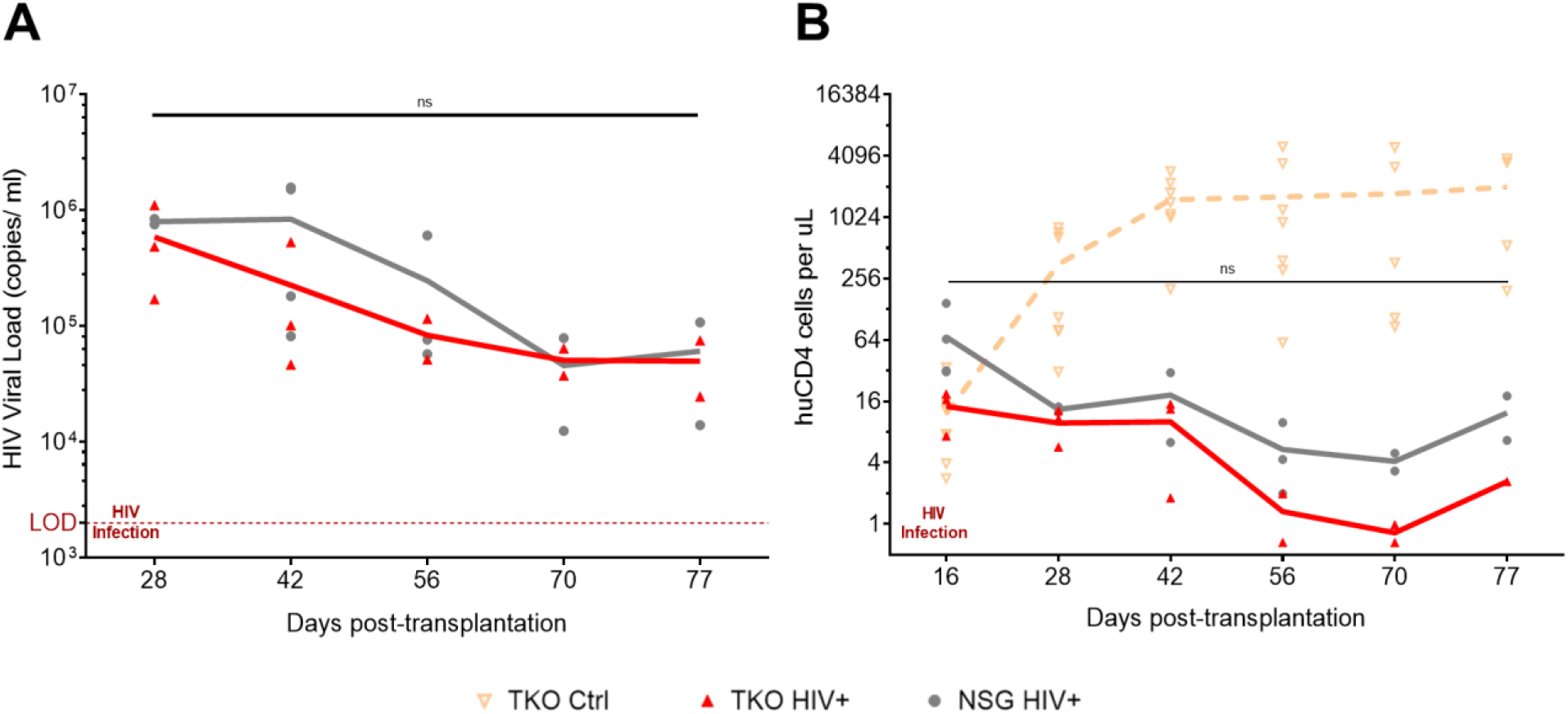
TKO hu-PBMC model supports HIV plasma viremia comparable to the NSG hu-PBMC model. TKO (n=3) and NSG (n=4) mice were transplanted with human PBMCs and 16 days later mice were infected with HIV-1_BaL_ (20 ng p24/mouse). Plasma HIV viremia and CD4^+^ T cells were monitored every 2 weeks by qRT-PCR and FACS analysis. As seen, the TKO hu-PBMC model supports HIV infection with robust plasma viremia (>2×10^3^ copies/ml) (**A**), and loss of CD4^+^ T cells over the 11 weeks, (**B**). CD4^+^ T cell counts from control TKO mice were added to compare HIV infected vs no HIV. Viremia in the TKO hu-PBMC model is comparable to the viremia seen in the NSG hu-PBMC model. (Note: mice were lost due to death at days 56 and 70). Statistical significance determined by mixed-effects analysis on Prism, with alpha = 0.05.

Next, we evaluated if HIV-1 infected TKO hu-PBMC mice would respond to oral combinatorial antiretroviral therapy (cART) similar to humans. In humans, cART medications block the replication of HIV leading to suppression of viral loads which in turn increases CD4^+^ T cell counts (15). To test the TKO hu-PBMC model, we treated mice orally for four weeks with cART composed of drugs that block new infections, without inhibiting viral production in infected cells. The cART regimen consisted of HIV nucleoside analog reverse transcriptase inhibitors Truvada® [tenofovir disoproxil fumarate (TDF) and emtricitabine (FTC)] (Gilead Sciences) and integrase inhibitor Isentress® [raltegravir (RAL)] (Merck), scaled down to the equivalent mouse dosage using the appropriate conversion factor (16). Medications were powered, mixed, and suspended in sweetened water solution (Medidrop® Sucralose, ClearH20) and provided in water bottles to mice as the sole water source. After two weeks of oral cART, plasma viral loads were significantly suppressed with some mice below the limit of detection (Fig. 4a). Plasma viremia was measured weekly while the mice were on oral cART for four weeks, in which all mice showed suppressed plasma viral loads. With suppressed plasma viremia, CD4^+^ T cell counts increased significantly while the mice received oral cART (Fig. 4b) modeling treatment in humans. When oral cART treatment was discontinued, there was a rebound of plasma viremia followed by a decline in CD4^+^ T cells (Fig. 4) modeling treatment in humans. Collectively, these results demonstrate the use of the TKO hu-PBMC model for testing antiviral drugs *in vivo*.

**Figure 4.**
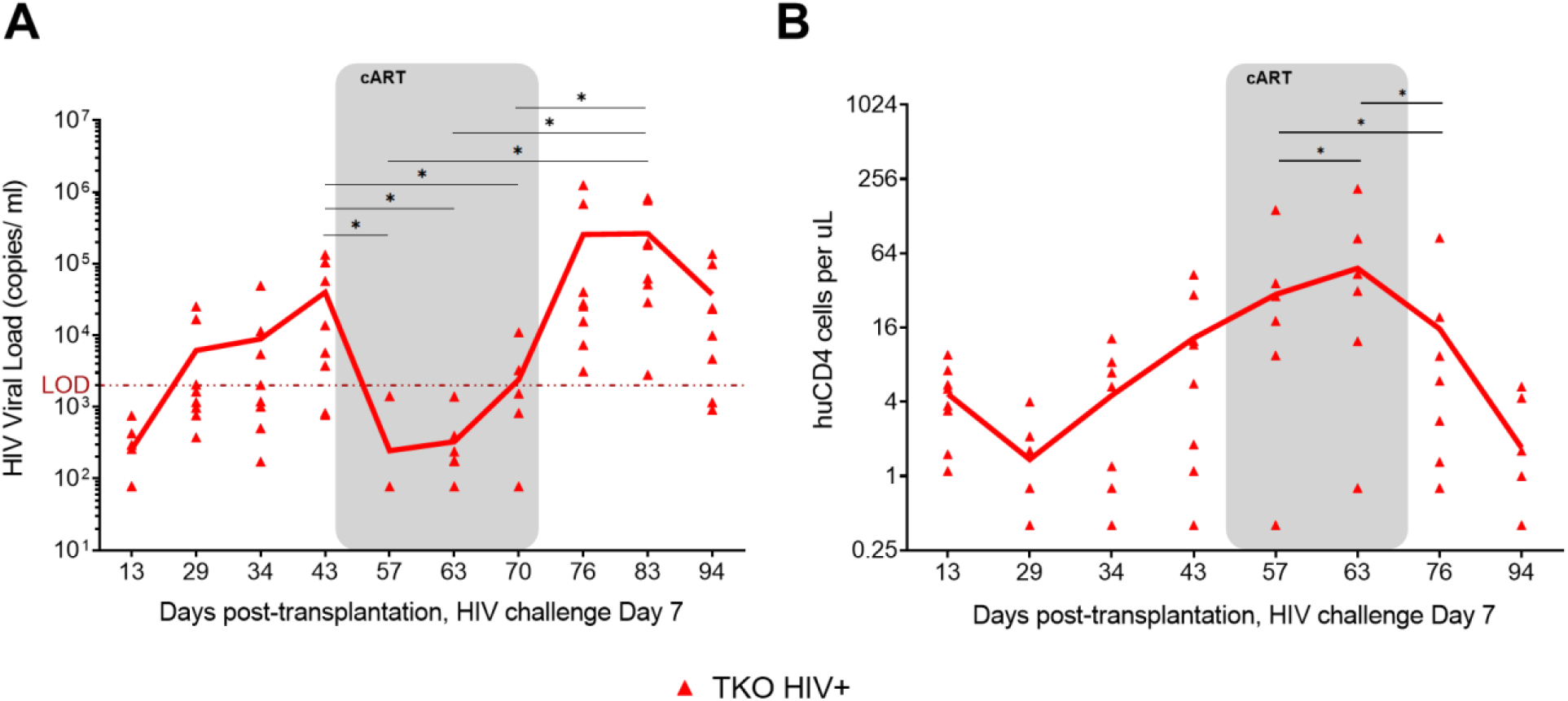
TKO hu-PBMC model response to oral cART with suppression of plasma viremia and CD4^+^ T cell protection. TKO hu-PBMC mice (n=8) were challenged with HIV-1BaL, 3 weeks after mice were given oral cART for 4 weeks. Two weeks after the start of oral cART plasma viremia was suppressed while oral cART was given and rebounded once oral cART was interrupted (**A**). A similar response was seen in CD4^+^ T cell protection while mice were on oral cART (**B**). The TKO model response to oral cART is similar to the NSG model (historical data). Statistical significance was determined by a one-tailed paired student’s t-test, with alpha = 0.05, *= p<0.05.

Lastly, we evaluated the HIV infection in the lymphoid tissues by immunohistochemistry. At the endpoint of the studies, tissues were collected and submitted for IHC staining as described earlier with the addition of HIV p24. There was positive staining for HIV p24 in bone marrow, liver, mesenteric lymph node, spleen, and rectum (Fig. 5). The positive staining of HIV p24 indicates that HIV-infected cells reside within the lymphoid tissues, including those deemed reservoirs in humans (bone marrow, lymph node, and spleen). While the TKO hu-PBMC model supports engraftment of CD4 T cells in the brain (Supplemental Fig. 1), HIV p24 staining in the brain was inconclusive (data not presented) indicating the need for further studies. Since the persistence of HIV despite ART is a critical factor in HIV cure strategies, we also wanted to evaluate the effect of oral cART on tissue reservoirs of infected mice. During oral cART, a cohort of mice with suppressed plasma viremia was humanely euthanized and tissues collected for IHC. In this group, there was a qualitative reduction of HIV p24 positive stained cells in the lymphoid tissue reservoirs of the bone marrow, lymph node, and spleen in mice on oral cART relative to untreated mice (Supplemental Fig. 3). This indicates that HIV infection persists in the TKO hu-PBMC model despite suppressive oral cART, demonstrating the use of the model to study HIV persistence and treatments.

**Figure 5.**
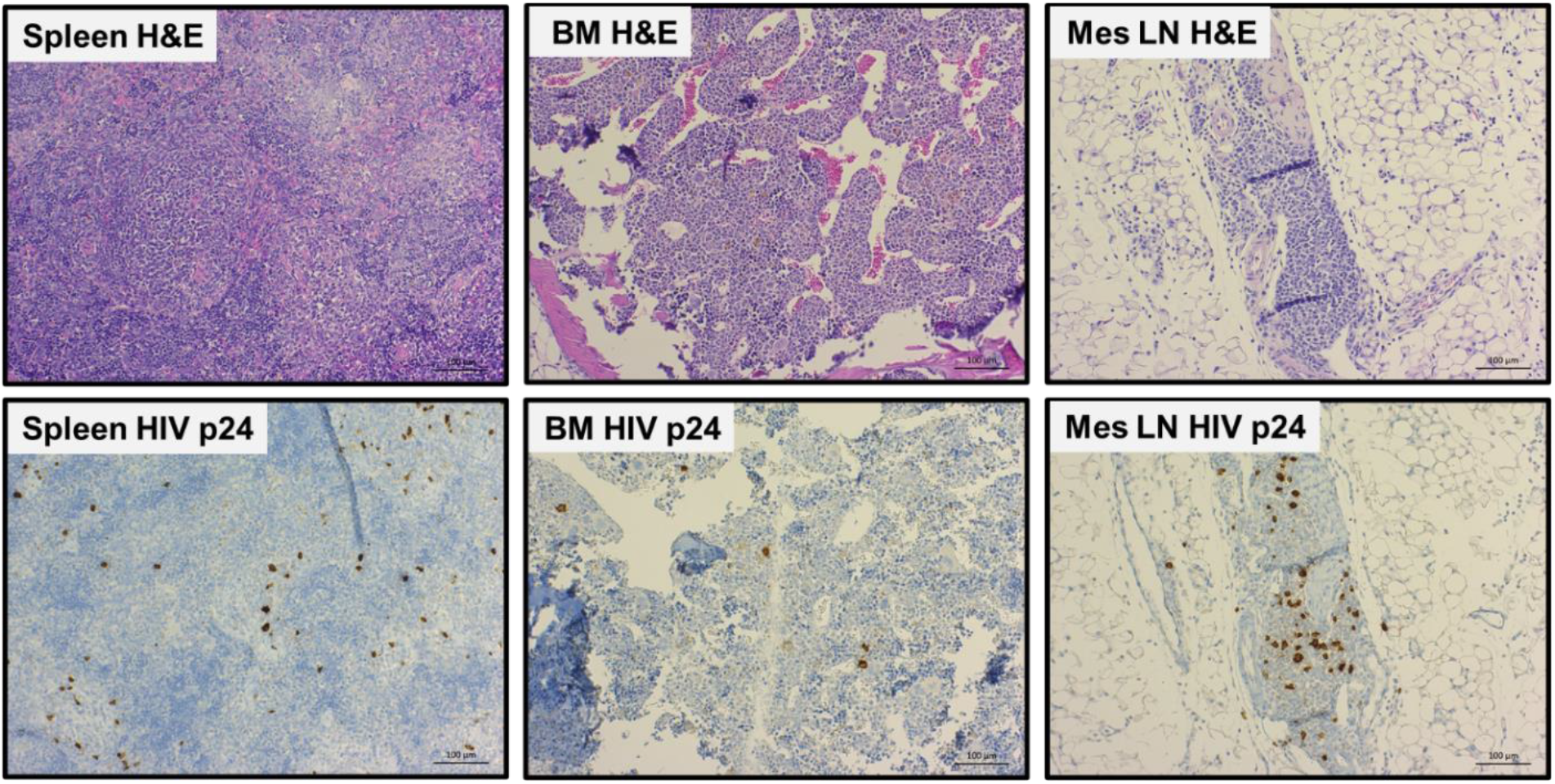
TKO hu-PBMC model supports HIV infection of lymphoid tissue reservoirs. TKO mice with robust and consistent peripheral CD45^+^ engraftment and robust plasma viremia were humanely euthanized and tissues (spleen, sternum (bone marrow), mesenteric lymph node) were collected for histology and IHC. Tissues were processed by the COH Solid Tumor Pathology Core for histology and immunohistochemistry: H&E, huCD3^+^, huCD4^+^, huCD8^+^, huCD14^+^/163^+^, and HIV p24. There is positive staining of HIV p24 (as seen above) indicating HIV infection in lymphoid tissue reservoirs, modeling HIV infection in humans. Representative images are shown.

### TKO hu-PBMC model has a delayed onset of GVHD compared to the NSG model

The major limitation with the NSG hu-PBMC model is the development of graft-versus-host disease (GVHD). To test whether the TKO hu-PBMC model would be resistant to the development of GVHD, clinical signs of graft-versus-host disease (GVHD) were monitored weekly after transplantation of mice and the date of the first sign of GVHD was recorded. Based on previous studies (10), we categorized GVHD as mild, moderate, or severe when observing clinical signs. Mild GVHD was defined as having a slightly ruffled hair coat with/out slightly hunched posture. Moderate GVHD was defined as having a ruffled hair coat with/out hunched posture, weight loss of 10-20%, and with/out alopecia. Severe GVHD was defined as having a ruffled hair coat, hunched posture, weight loss greater than 20%, and with/out alopecia and lethargy. The median onset of mild GVHD clinical signs in NSG mice was 28 days, while in TKO mice it was 52 days (Fig. 6a). The TKO hu-PBMC model has a delayed onset of mild GVHD of 24 days compared to the NSG model. When we monitored for moderate GVHD, the median onset moderate GVHD clinical signs in NSG mice was 50.5 days, while in TKO mice it was 66 days (Fig. 6b). The TKO hu-PBMC model has a delayed onset of moderate GVHD of 15.5 days compared to the NSG model. Finally, when we monitored for severe GVHD, the TKO hu-PBMC model has a delayed onset of 18 days compared to the NSG model (data not shown). These results show that the TKO hu-PBMC model has a delayed onset of mild, moderate, and severe GVHD compared to the NSG hu-PBMC model.

**Figure 6.**
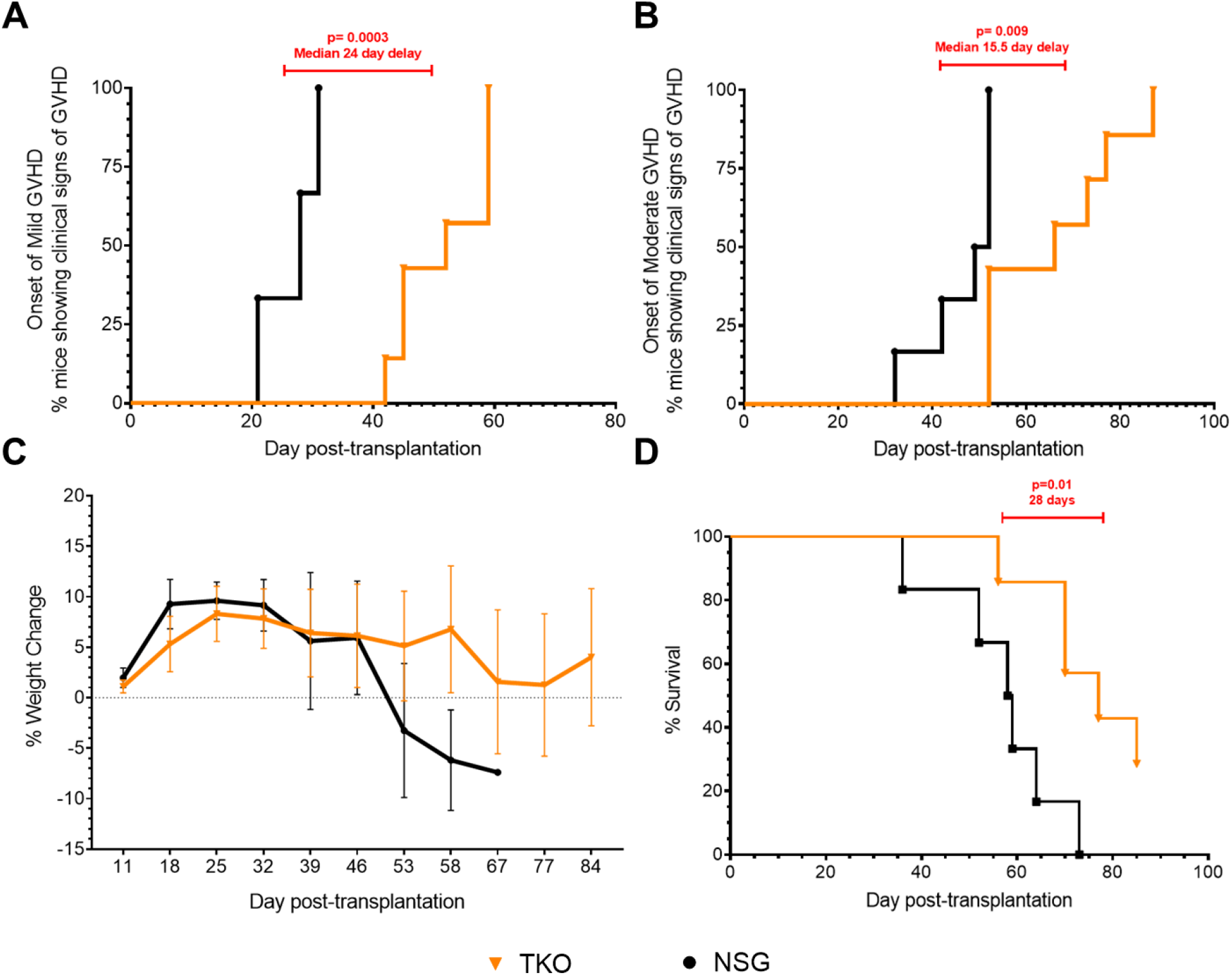
Delayed onset of GVHD in the TKO model compared to the NSG model. TKO (n=7) and NSG (n=6) hu-PBMC mice were monitored twice weekly for clinical signs of GVHD. Mild GVHD was defined as having a slightly ruffled hair coat with/out slightly hunched posture. Moderate GVHD was defined as having a ruffled hair coat with/out hunched posture, weight loss of 10-20%, and with/out alopecia. TKO mice had a 24-day delayed onset of mild GVHD (**A**), and a 15.5 delay onset of moderate GVHD compared to NSG mice (**B**). Monitoring weight weekly, TKO mice maintained and lost less weight percentage compared to NSG mice (**C**). Evaluating survival time, TKO mice had a 28-day longer survival time compared to NSG mice (**D**). Statistical significance determined by Log-rank (Mantel-Cox) test on Prism, with alpha = 0.05.

Weight loss is a clinical sign of GVHD and a major concern for the welfare of the mice (17). We monitored the weight of the mice weekly to measure percent weight loss over time which revealed that NSG mice had greater and faster weight loss compared to TKO mice (Fig. 6c). TKO mice were able to maintain their weight over the length of the study, while NSG mice began to lose weight after day 46. Coupled with the weight, we also monitored the survival time of the mice. The date of death was defined as the day mice had to be euthanized for humane endpoints or the day mice were found dead. TKO mice survived an average of 26 days longer than NSG mice over the length of the study (Fig. 6d). All NSG mice were dead by day 73, while two TKO mice survived until the end of the study, day 85. These results show that the TKO hu-PBMC model maintains weight and has a longer survival time compared to the NSG hu-PBMC model.

To evaluate whether there were differences in plasma inflammatory cytokines produced by the TKO and NSG hu-PBMC models, a human inflammatory cytokine assay was performed on the plasma of the mice collected throughout the study. The samples were tested for Human IL-1α, IL-1β, IL-2, IL-4, IL-5, IL-6, IL-10, IL-12, IL-13, IL-15, IL-17, IL-23, IFNγ, TNFα, and TNFβ using on Quansys Biosciences’ Q-Plex™ Human High Sensitivity multiplexed ELISA array. While there were no significant differences in most cytokines assayed (Supplemental Fig. 4), the NSG hu-PBMC mice had significantly more plasma TNFβ compared to the TKO mice (Fig. 7). The low plasma levels of TNFβ in the TKO hu-PBMC mice may contribute to the delayed onset of GVHD compared to the NSG model as further discussed later.

**Figure 7.**
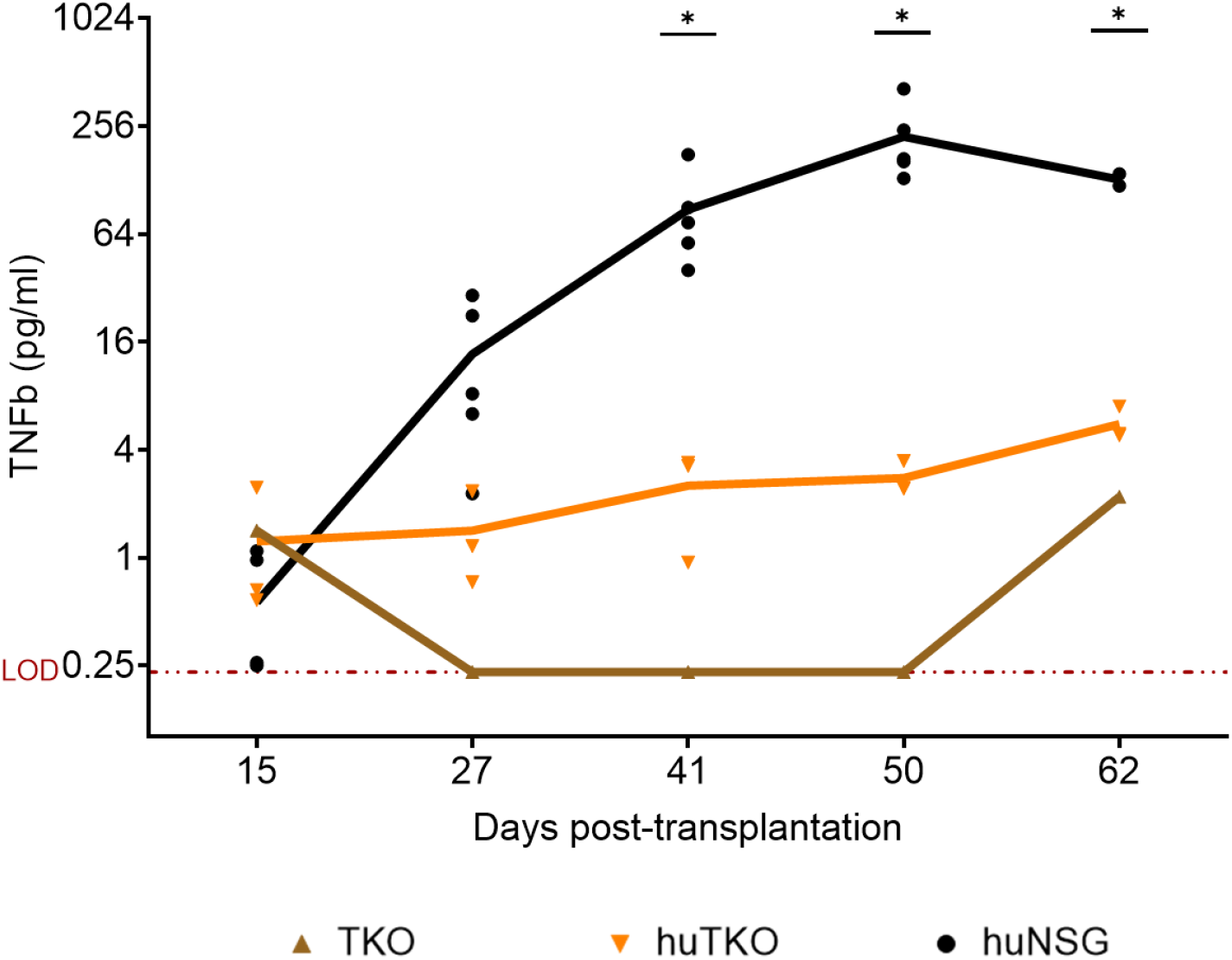
NSG hu-PBMC mice have more circulating plasma TNFβ compared to TKO hu-PBMC mice. Plasma samples were collected every 2 weeks and submitted to Quansys Biosciences (Logan, UT) to be tested on Quansys Biosciences’ Q-PlexTM Human High Sensitivity multiplexed ELISA array. All other human inflammatory cytokines had no difference between TKO (n=3) and NSG hu-PBMC (n=5) mice, except for TNFβ. NSG hu-PBMC mice have more circulating plasma TNFβ compared to TKO hu-PBMC mice. Statistical significance determined by mixed-effects analysis on Prism, with alpha = 0.05; p values= *<0.05.

## Discussion

TKO mice have previously been shown to be GVHD resistant when used to develop a hu-BLT mouse model (12); however, that model requires human fetal tissues and other practical limitations including its reproducibility and availability to the research community (18). In this study, we developed a novel hu-PBMC mouse model that supports HIV-1 infection and is more resistant to graft-versus-host disease (GVHD), than the previously established NSH hu-PBMC model (7).

We observed consistent engraftment of human CD45^+^ leukocytes and CD3^+^ lymphocytes in TKO mice after intraperitoneal transplantation of human PBMCs (Fig. 1). Further analysis of T cell subpopulations showed the robust circulation of CD4^+^ and CD8^+^ T cells in the peripheral blood of the TKO hu-PBMC model (Fig. 1). When comparing the model to the commonly used NSG hu-PBMC model, we observed no significant difference in engraftment of human cells between the models. Immunohistochemistry analysis of the tissues showed that the TKO hu-PBMC model supports the engraftment and development of lymphoid tissues including bone marrow, lymph node, and spleen with human T cells (Fig. 2, Table 1, and Supplemental Fig. 1). The presence of human T cells in the lymphoid tissues is notable because of the critical role T cells play in the establishment and maintenance of persistent HIV-1 infection in humans.

**Table 1.**
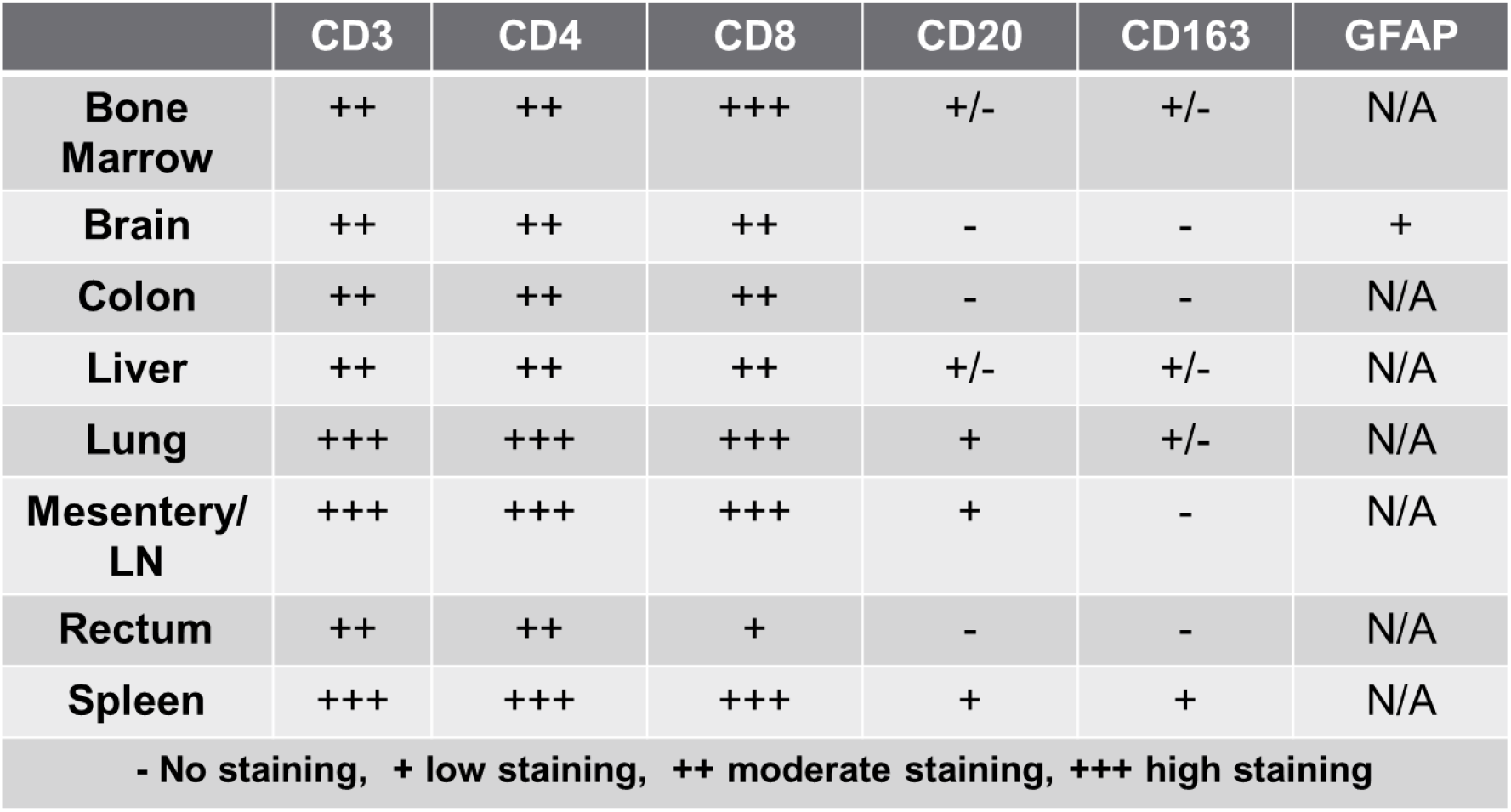
Chart of engraftment of human immune cells in different tissues.

To test the utility of the novel TKO hu-PBMC model, we challenge mice with HIV-1BaL virus intraperitoneally, rectally, and vaginally. Challenging via all three routes produced robust plasma viremia and depletion of CD4^+^ T cells over time (Fig. 3 and Supplemental Fig. 2) modeling human pathogenesis. When comparing the TKO and NSG models, there were no significant differences in plasma viremia or CD4^+^ T cell depletion. We also observed robust HIV p24 antibody staining in bone marrow, lymph node, and spleen when we analyzed lymphoid tissue by IHC (Fig. 5). When we placed the HIV-infected TKO hu-PBMC mice on oral cART for four weeks, we observed a suppression of plasma viral viremia and an increase in CD4^+^ T cells (Fig. 4). Once oral cART was stopped, there was a rebound of plasma viremia and CD4^+^ T cell depletion as seen in humans.

Lastly, we observed resistance to the development of GVHD in the TKO hu-PBMC model. Weekly observation for GVHD clinical signs, revealed a median 24-day delay on the onset of GVHD and a median 15.5-day delay on the onset of GVHD in the TKO model compared to the NSG model (Fig. 6). We also observed a greater ability of the TKO hu-PBMC mice to maintain their body weights compared to the NSG mice over the length of the study. Survival time was increased in the TKO model compared to the NSG model, with TKO hu-PBMC mice having an average 26-day longer survival time (Fig. 6). We observed significantly lower levels of plasma TNFβ in the TKO hu-PBMC mice compared to the NSG mice (Fig. 7). TNFβ, also known as lymphotoxin alpha (LTα), is a TNF-superfamily member and plays a crucial role in the development and orchestration of robust immune responses (19). Activated CD4^+^ Th subsets Th1 and Th17, but not Th2, express surface LTα, as do CD8^+^ T cells and B cells (19, 20). LTα has been previously shown to be an important contributor to GVHD pathogenesis (21). Chiang et al. 2012, described the use of a humanized anti-LTα monoclonal antibody (MLTA3698A) that specifically depleted activated LT-expressing human donor T and B cells, resulting in prolonged survival times in the hu-SCID mouse model of GVHD (19). Based on these findings, it may be possible that the low production of TNFβ in the TKO hu-PBMC model may contribute to the delayed onset of GVHD compared to the NSG model. However, further studies are warranted to study the role of TNFβ in the TKO model.

As shown, our novel TKO hu-PBMC model supports and models HIV-1 infection and has resistance to GVHD development when compared to the NSG hu-PBMC model. The delayed onset of GVHD and increased survival time of the TKO hu-PBMC mice make them a valuable tool for studying HIV-1 infection, pathogenesis, and evaluating new therapies.

## Methods

### Mice

NOD.Cg-*Prkdc*^*scid*^ *Il2rg*^*tm1Wjl*^/SzJ (NSG) mice (JAX stock #005557) and B6.129S-*Rag2*^*tm1Fwa*^ *CD47*^*tm1Fpl*^ *Il2rg*^*tm1Wjl*^/J (TKO) mice (JAX stock #025730) were purchased from the Jackson Laboratory (Bar Harbor, ME). Mice were maintained in accordance with the Guide for the Care and Use of Laboratory Animals and were housed in SPF conditions. All experiments were performed according to the guidelines of the Institutional Animal Committee of the Beckman Research Institute of The City of Hope, IACUC protocol #16095. Experiments were done in accordance with Public Health Service Office of Laboratory Animal Welfare regulations. Beckman Research Institute of The City of Hope is fully accredited by the Association for Assessment and Accreditation of Laboratory Animal Care.

### Preparation and engraftment of PBMCs

Human PBMCs were obtained from peripheral venous blood of healthy donors from the Apheresis Center at the City of Hope, in accordance with the COH Institutional Review Board protocol # 17155. Human PBMCs (1×10^7^ cells), purified by density gradient centrifugation using SepMate™ tubes and Lymphoprep™ media (Stem Cell Technologies, Vancouver, BC), were resuspended in sterile phosphate-buffered saline (PBS) and injected intraperitoneally (IP) into NSG mice.

### HIV-1 infection and oral cART therapy

Once engraftment of mice was confirmed, they were challenged with HIV-1_BaL_virus (20 ng p24/mouse) by IP injection while under general isoflurane anesthesia. HIV-1_BaL_ cell-free virus was obtained from the NIH HIV Reagent Program (HIV-1_BaL_, ARP-510) then cultured, titered, and frozen. For the rectal and vaginal challenge, mice were anesthetized with isoflurane to reach a deep surgical plane as previously described (22, 23). A pipet was rubbed in the genital area, to stimulate the emptying of the rectum. For the rectal challenge, mice were placed in ventral recumbency, and the tail was pulled up to expose the rectum. 10-20 µL of HIV-1_Bal_ virus (4-8 ng p24) was atraumatically pipetted into the rectum. For the vaginal challenge, mice were placed in dorsal recumbency, and the tail was pulled and wrapped around fingers to expose the vagina. 10-20 µL of HIV-1_Bal_ virus (4-8 ng p24) was pipetted atraumatically into the vagina. Mice were held with hindlimbs elevated for 3-5 min, then transferred to the home cage to recover in a position with hindlimbs elevated. All procedures involving HIV were performed in biosafety level 2+ (BSL2+) conditions in accordance with protocols approved by the institutional biosafety committee of the City of Hope. Mice were bled by retro-orbital bleeding, and peripheral blood cell populations and plasma viral loads were analyzed periodically using flow cytometry and qRT-PCR.

Infected mice with demonstrable viral infection were treated orally for four-to-six weeks with cART composed of drugs that block new infections, without inhibiting viral production in infected cells. The cART regimen consisting of Truvada® [tenofovir disoproxil fumarate (TDF; 300 mg/tablet), emtricitabine (FTC; 200 mg/tablet) (Gilead Sciences)] and Isentress® [raltegravir (RAL; 400 mg/tablet) (Merck)], scaled down to the equivalent mouse dosage using the appropriate conversion factor, was administered in a drinking water formulation (sweetened water gel, Medidrop® Sucralose, ClearH20). For 400 mL Medidrop®, ½ Truvada tablet, and ½ Isentress tablet were crushed to powder and mixed by shaking the bottle to a homogenous solution; medicated water was changed weekly. Doses of cART drugs were calculated based on previous studies using the same delivery system(16). Mice were bled by retro-orbital bleeding, and peripheral blood cell populations and plasma viral loads were analyzed periodically using flow cytometry and qRT-PCR. Mice were bled weekly for qPCR, while FACS analysis was sometimes done every two weeks due to limited sample volumes. Mice that had suppressed or undetectable plasma viral loads were followed for 4-6 weeks, at which time, the cART regimen was withdrawn. The mice were bled and assayed for the rebound of plasma viremia and peripheral cell composition periodically until euthanized.

### Flow cytometry

Samples of peripheral blood were collected by retro-orbital bleeding under general anesthesia and stained for 30min with BV711-conjugated antihuman CD3, APC-conjugated antihuman CD4, BB515-conjugated antihuman CD8, and BUV395-conjugated antihuman CD45 (all from BD Biosciences, San Jose, CA). Stained peripheral blood samples were then lysed with red blood cell lysis buffer and absolute cell counts calculated using BD Liquid Counting Beads (BD Biosciences, San Jose, CA). Flow cytometry was performed using BD Fortessa II instrument (BD Biosciences) and analyzed with FlowJo software.

### Intracellular HIV p24 Flow cytometry

Samples of peripheral blood and single-cell suspensions of mouse spleen and bone marrow (femurs +/-tibias) were collected at the time of euthanasia. Single-cell suspensions were made following previously established protocols (24). Briefly, for bone marrow cells, femurs and tibia were dissected and collected from euthanized mice and placed in ice-cold PBS. Bones were cleaned thoroughly to remove all connective and muscle tissue, then using a scalpel blade the heads of the bones were removed. Bones were placed in a 0.5ml microcentrifuge tube with a premade hole by using a 20g needle. Bones were placed cut surface down in 0.5ml tubes and those tubes were placed in 1.5ml microcentrifuge tubes and centrifuged at >10,000 x g for 15 sec. The cell pellet was resuspended in ACK lysis buffer incubated for 5 min and washed with PBS. Cells were resuspended in PBS+2% FBS and then processed for FACS staining or frozen in 10% Cryostor (Stem Cell Technologies, Vancouver, BC).

For spleen cells, spleens were collected from euthanized mice and placed in ice-cold PBS. Spleens were processed by placing them on a 40 µm cell strainer and using a syringe plunger to mash the tissue through the strainer and washed with PBS. Cells were centrifuged at 400 x g for 10 min and then the cell pellet was resuspended in ACK lysis buffer incubated for 5 min and washed with PBS. Cells were resuspended in PBS+2% FBS and then processed for FACS staining or frozen in 10% Cryostor (Stem Cell Technologies, Vancouver, BC). For intracellular staining, BD Cytofix/Cytoperm™ kit (BD Biosciences, San Jose, CA) was used following the manufacture’s protocol. After surface markers staining (CD45^+^, CD3^+^, CD4^+^, CD8^+^), cells were permeabilized and intracellular staining of RD1-conjugated antiHIV-1 p24 (Beckman Coulter, Brea, CA). Flow cytometry was performed using BD Fortessa II instrument (BD Biosciences) and analyzed with FlowJo software.

### Plasma HIV qRT-PCR

Plasma viremia was assayed using one-step reverse transcriptase real-time PCR [TaqMan assay] with an automated CFX96 Touch™ Real-Time PCR Detection System (Bio-Rad). qPCR primer sets were taken from previously published studies(16). HIV-1 level in peripheral blood was determined by extracting RNA from blood plasma using the QIAamp Viral RNA mini kit (Qiagen) and performing Taqman qPCR using either a primer and probe set targeting the HIV-1 LTR region [FPrimer: GCCTCAATAAAGCTTGCCTTGA, RPrimer: GGCGCCACTGCTAGAGATTTT, Probe: 5’FAM/AAGTAGTGTGTGCCCGTCTGTTGTGTGACT /3IABkFQ] or the HIV-1 Pol region [FPrimer: GACTGTAGTCCAGGAATATG, RPrimer: TGTTTCCTGCCCTGTCTC, Probe: 5’Cy5/CTTGGTAGCAGTTCATGTAGCCAG/3’IABkFQ], using the TaqMan Fast Virus 1-Step Master Mix (Applied Biosystems). According to the manufacturer’s instruction (QIAamp Viral RNA mini kit (Qiagen)), the protocol is designed for purification of viral RNA from minimal 140 μL plasma. In a standard Taqman qPCR-based HIV-1 plasma viral load test, the limit of detection (LOD) is typically about 40 copies/ml when viral RNA isolated from 140 μL of plasma sample is applied. In our animal study, the plasma sample was expanded by dilution (generally 1 to 3 dilution) because a limited volume of plasma (20-60 μL) was available. The LOD of the diluted samples was around ∼2000 RNA copies/ml using the HIV LTR primer and ∼500 RNA copies/ml using the HIV Pol primer under our experimental condition. Therefore, we defined that the value below those LOD numbers is undetectable.

### Immunohistochemistry

Samples of skin, lung, liver, spleen, small and large intestine, sternum bone marrow, reproductive tract, and brain tissue from mice were examined at the time of euthanasia. Tissues were submitted to the Pathology Solid Tumor Core of Beckman Research Institute at the City of Hope. These samples were fixed with 10% formaldehyde, embedded in paraffin, and stained with hematoxylin and eosin (H&E). H&E slides were analyzed for histological signs of graft-versus-host disease.

Immunohistochemical staining was performed on 5-μm thick sections from PFA-fixed, paraffin-embedded tissue using a standard protocol. Primary antibodies used for immunohistochemistry: rabbit antihuman-CD3^+^ (SGV6 clone, Ventana), rabbit antihuman-CD4^+^ (SP35 clone, Ventana), rabbit antihuman-CD8^+^ (SP57 clone, Ventana), mouse antihuman-CD20^+^ (L26 clone, Ventana), mouse antihuman-CD163^+^ (MRQ-42 clone, Ventana), mouse antihuman-CD68^+^ (KP-1 clone, Ventana), mouse antihuman-GFAP (GA5 clone, BOND RTU), and mouse anti-HIV p24 (Kal-1 clone, Dako). DAB staining was performed using a Ventana Ultraview DAB detection kit in a Ventana BenchMark XT processor (Ventana Medical Systems, a division of Roche). In addition, staining of a non-humanized mouse with the same primary antibodies was used as an additional control.

### Monitor Sign of Graft-versus-Host Disease

Clinical signs of graft-versus-host disease (GVHD) were monitored weekly after transplantation of mice and the date of the first sign of GVHD was recorded. Based on previous studies (10), clinical signs observed and reported were: slightly ruffled hair coat, ruffled hair coat, slightly hunched posture, hunched posture, lethargy, alopecia, and weight loss ranging from 10-20% and greater than 20%. We categorized GVHD as mild, moderate, or severe when observing clinical signs. Mild GVHD was defined as having a slightly ruffled hair coat with/out slightly hunched posture. Moderate GVHD was defined as having a ruffled hair coat with/out hunched posture, weight loss of 10-20%, and with/out alopecia. Severe GVHD was defined as having a ruffled hair coat, hunched posture, weight loss greater than 20%, and with/out alopecia and lethargy. Lab members with animal husbandry/care experience were in charge of the weekly observations. The onset of mild, moderate, and severe GVHD clinical signs was recorded. When mice presented with severe GVHD clinical signs with extreme lethargy or moribund, they were humanely euthanized, and the date was recorded. For Meir-Kaplan survival curves, date of death either by humane euthanasia or found dead were used.

### Human cytokine multiplexed ELISA assay

Plasma samples from mice (40-100 µL) were saved and frozen in −80C at every blood collection time point of the study. Samples were shipped to Quansys Biosciences (Logan, UT) to be tested on Quansys Biosciences’ Q-Plex™ Human High Sensitivity multiplexed ELISA array. The samples were tested for Human IL-1α, IL-1β, IL-2, IL-4, IL-5, IL-6, IL-10, IL-12, IL-13, IL-15, IL-17, IL-23, IFNy, TNFα, and TNFβ. Samples were thawed at ambient temperature and then kept cold on ice. Thawed samples were then diluted with the appropriate Quansys sample dilution buffer. Samples were diluted at ratios (sample:total volume) of 1:5 (20%), 1:20 (5%), and 1:100 (1%). Polypropylene low-binding 96-well plates were used to prepare the samples and standards prior to loading the Q-Plex TM plate. Each dilution was measured in duplicate for a total of 6 wells per sample. Antigen standard curves were performed in duplicate diluting the antigen standard 1:3 for 10 points with two negative points. The sample and antigen standard incubation was extended from one hour to two hours, and the detection of secondary antibody incubation was extended from one hour to two hours. An image with a 270 second exposure time was captured using a Q-View™ Imager LS and Q-View™ Software. Levels of luminescent units or pixel intensity units were then measured by the Q-View™ Software. The duplicate standard curves are fit by the Q-View™ Software which allows for the selection of multiple non-linear and linear equations to fit the standard curve. Optimal curve fits are determined automatically by the software by evaluating the recovery of the calibrator standards.

### Statistical analysis

Unless otherwise noted, error bars in all figures, both in the main part of the paper, represent the standard error of the mean (SEM). GraphPad Prism software was used for statistical analyses: Student’s t-test, ANOVA, mixed-effects analysis, or Log-rank statistical tests were used, and differences were considered statistically significant when p < 0.05.

## Supporting information

all supplemental figures

## Acknowledgments

The following reagent was obtained through the NIH HIV Reagent Program, Division of AIDS, NIH: Human Immunodeficiency Virus-1 Ba-L, ARP-510, contributed by Dr. Suzanne Gartner, Dr. Mikulas Popovic, and Dr. Robert Gallo. We thank the City of Hope core facilities that were used in this study: Analytical Cytometry Core (ACC), Center for Comparative Medicine (CCM) and its breeding core, and the Pathology Research Services supported by the National Cancer Institute of the National Institutes of Health under award number P30CA33572. We thank Dr. Kevin Morris and Dr. Xiuli Wang for funding support of this project. We thank members of the Burnett Lab for discussions and suggestions.

## Author Contributions

L.H. and J.C.B. conceived and designed the study and wrote the manuscript. L.H. and L.E. designed and performed the experiments. L.H. curated the data.

