## Supplementary material for "Novel GVHD resistant humanized-PBMC mouse model for preclinical HIV research": all supplemental figures

### Supplemental Material

Supplemental Figure 1- Representative IHC images for TKO hu-PBMC mice

Supplemental Figure 2- Rectal and Vaginal HIV challenge

Supplemental Figure 3- IHC images for HIV mice and while on oral cART

Supplemental Figure 4- Plasma Cytokine Assay

### Supplemental Fig. 1- Representative IHC images for TKO hu-PBMC mice

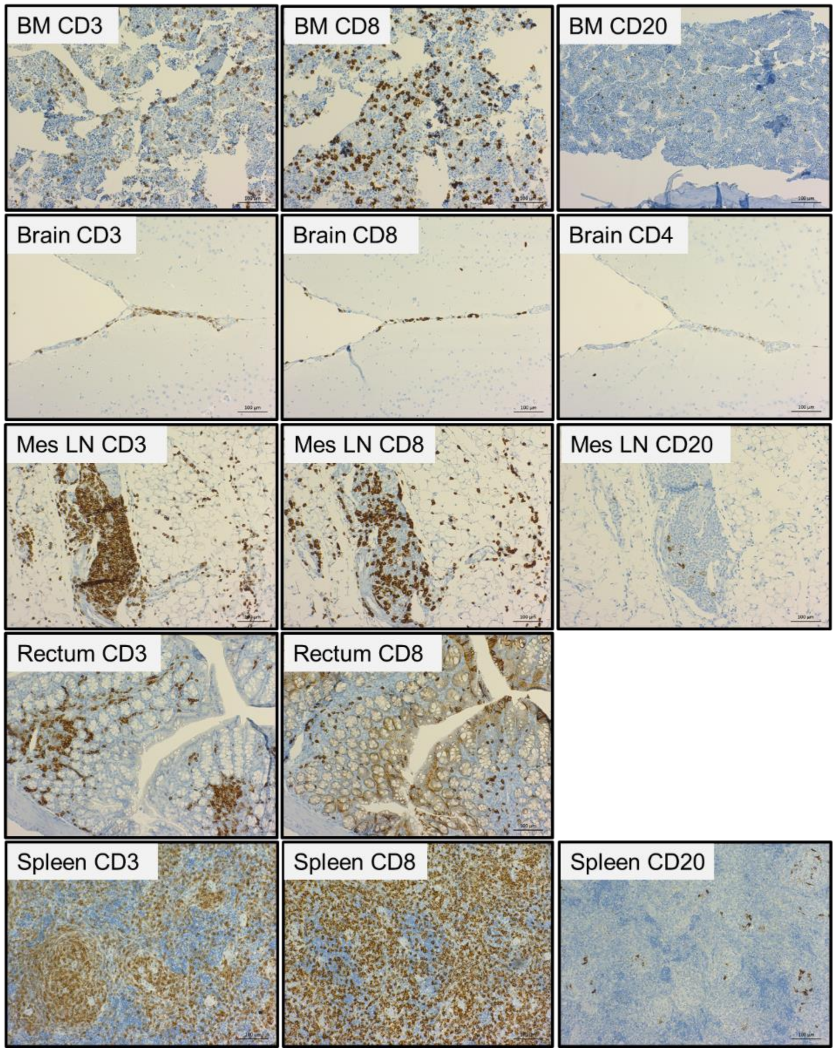


***Supplemental Figure 1. TKO hu-PBMC model supports the engraftment of multiple human hematopoietic cells in various tissues***. TKO mice with robust and consistent peripheral $CD45^+$ engraftment were humanely euthanized, and tissues were collected for histology and IHC. Tissues were processed by the COH Solid Tumor Pathology Core for histology and immunohistochemistry: H&E, $huCD3^+$, $huCD4^+$, $huCD8^+$, $huCD14^+/163^+$, $huCD20^+$, $huCD56^+$, huGFAP. Representative images are shown. Table 1 indicates the intensity of cellular markers staining in different tissues.

### Supplemental Fig. 2- Rectal and Vaginal HIV challenge

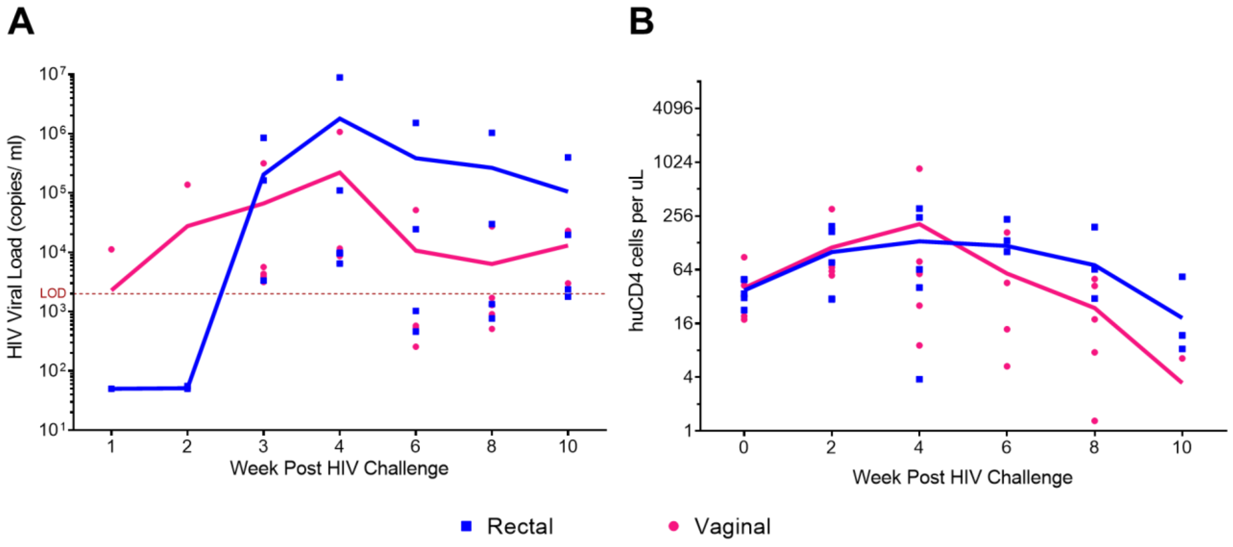


***Supplemental Figure 2. TKO hu-PBMC mice model HIV infection when challenged via rectal and vaginal routes.*** TKO mice were transplanted with human PBMCs and 16 days later mice were infected with HIV-1$_{BaL}$ (20 ng p24/mouse). Male mice (n=5) were challenged rectally while female mice (n=5) vaginally. Plasma HIV viremia and CD4$^+$ T cells were monitored every 2 weeks by qRT-PCR and FACS analysis. As seen, the TKO hu-PBMC model supports HIV infection with robust plasma viremia (>2x10$^3$ copies/ ml) (**A**), and loss of CD4$^+$ T cells over the 10 weeks, (**B**). (Note: mice were lost due to death at weeks 4 and 8).

**Supplemental Fig. 3- IHC images for HIV mice and while on oral cART**

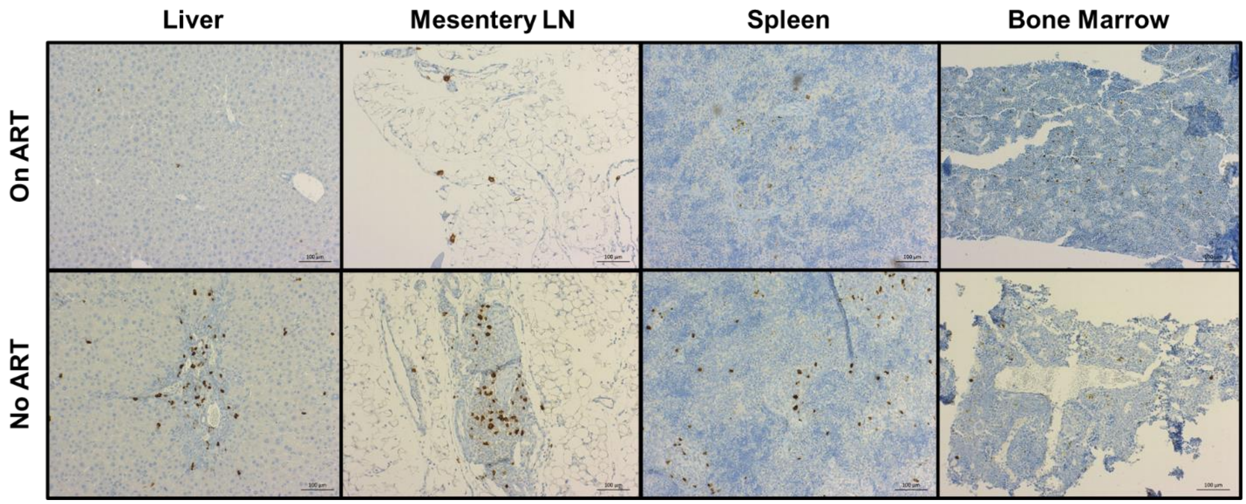


***Supplemental Figure 3. There is a qualitative reduction in HIV p24 staining when TKO hu-PBMC HIV+ mice receive oral cART compared to no oral cART.*** TKO hu-PBMC mice were challenged with HIV-$1_{BaL}$, 3 weeks after mice were given oral cART for 4 weeks. A cohort of mice was euthanized while on oral cART and IHC analysis was done on lymphoid tissues. Tissues were processed by the COH Solid Tumor Pathology Core for histology and immunohistochemistry: H&E, $huCD3^+$, $huCD4^+$, $huCD8^+$, and HIV p24. There is a qualitative reduction in HIV p24 positive staining for mice receiving oral cART compared to mice not receiving oral cART, modeling persistent HIV infection in humans. Representative images are shown.

### Supplemental Fig. 4- Plasma Cytokine Assay

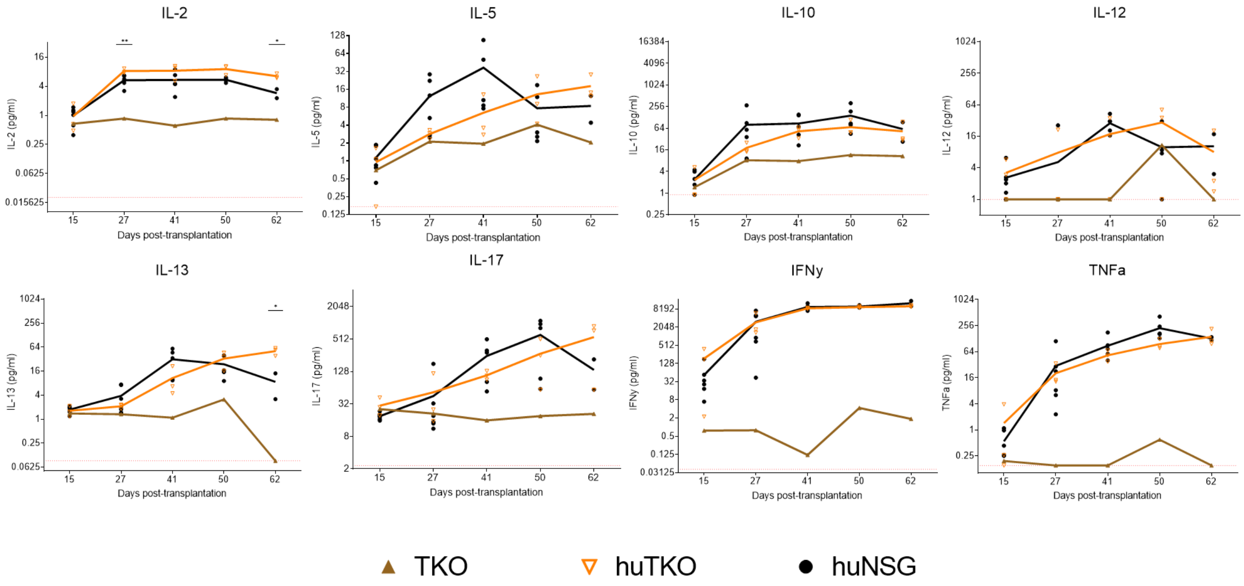


***Supplemental Figure 4. Plasma cytokine assay for TKO and NSG hu-PBMC models.*** Plasma samples were collected every 2 weeks throughout the study and frozen in -80C for assaying later. Plasma samples were submitted to Quansys Biosciences (Logan, UT) to be tested on Quansys Biosciences' Q-PlexTM Human High Sensitivity multiplexed ELISA array. The samples were tested for Human IL-1α, IL-1β, IL-2, IL-4, IL-5, IL-6, IL-10, IL-12, IL-13, IL-15, IL-17, IL-23, IFNγ, TNFα, and TNFβ. All other human inflammatory cytokines had no difference between TKO and NSG hu-PBMC mice, except for TNFβ. Statistical significance determined by mixed-effects analysis on Prism, with alpha = 0.05; p values= *<0.05, **<0.005.
